# Electrophysiological correlates of semantic dissimilarity reflect the comprehension of natural, narrative speech

**DOI:** 10.1101/193201

**Authors:** Michael P. Broderick, Andrew J. Anderson, Giovanni M. Di Liberto, Michael J. Crosse, Edmund C. Lalor

## Abstract

Understanding natural speech requires that the human brain convert complex spectrotemporal patterns of acoustic input into meaning in a rapid manner that is reasonably tightly time-locked to the incoming speech signal. However, neural evidence for such a time-locked process has been lacking. Here, we sought such evidence by using a computational model to quantify the meaning carried by each word based on how semantically dissimilar it was to its preceding context and then regressing this quantity against electroencephalographic (EEG) data recorded from subjects as they listened to narrative speech. This produced a prominent negativity at a time-lag of 200– 600 ms on centro-parietal EEG electrodes. Subsequent EEG experiments involving time-reversed speech, cocktail party attention and audiovisual speech-in-noise demonstrated that this response was exquisitely sensitive to whether or not subjects were understanding the speech they heard. These findings demonstrate that, when successfully comprehending natural speech, the human brain encodes meaning as a function of the amount of new information carried by each word in a relatively time-locked fashion.

---

In everyday life, people routinely process heard speech at rates in the range of 120 to 200 words per minute^1,2^. Unlike in the case of reading, listeners typically do not have much control over the rate at which these words are presented and they usually cannot replay the presentation of those words. Thus, successful speech comprehension must involve efficient, online mechanisms in the brain whereby each word is processed in a relatively time-locked fashion. In addition, it is well established that the processing of words does not happen in isolation, but is strongly influenced by the surrounding conversational context^3^. The field of psycholinguistics has long been concerned with how context rapidly impacts upon word processing to facilitate speech comprehension^4,5^. More recently interest has been placed on quantitatively modeling the effects of different aspects of context. This has included syntax^6^, and, following the introduction of large scale models for representing the contextual-usage meaning of words^7^, also semantics^8^, or both^9^.

While these models – and the experiments on which they are based – have greatly deepened our understanding of psycholinguistics, there has been a marked lack of electrophysiological evidence for the time-locked processing of meaning that must underpin natural speech comprehension. This is a shame as an electrophysiological index of such processing would be of great benefit for arbitrating between different psycholinguistic models, and could have important implications for research on language processing in numerous cohorts. Valuable insights into the semantic processing of speech have been provided by the well-known N400 component of the event-related potential^10^. However, the N400 literature has been dominated by paradigms focused on single, usually incongruous, words within specially constructed sentences, and has had much less to say about how ongoing neural activity reflects the computations that underpin natural, narrative speech comprehension. Furthermore, the use of the classic N400 paradigm has made it difficult to fully understand how selective attention and variations in intelligibility affect the semantic processing of speech under naturalistic conditions.

Here, in an attempt to address these issues, we build on the relatively recent discovery that the dynamics of cortical activity “track” the dynamics of natural, ongoing speech^11–13^. Much of this work has focused on how electrophysiological signals entrain to the dynamics of the speech envelope^12,13^, with measures of this entrainment having been shown to be affected by attention^14,15^ and intelligibility^16^. And, more recently, there have been efforts to more directly link this neural tracking to the processing of speech at different hierarchical levels^17^, including at the level of phonemes and phonetic features^18^. However, to date, no work has shown that this ongoing electrophysiological activity reflects anything about the semantic processing of natural speech, and how that is affected by attention and intelligibility. This is the goal of the present work.

## RESULTS

We acquired electroencephalographic (EEG) data from subjects as they listened to narrative speech in the form of audiobook recordings. To relate the neural data to the semantic processing of this speech, we first wished to parameterize the speech stimuli such that individual words were quantified according to their semantic context. There are many ways to do this. Inspired by the brain’s sensitivity to incongruous new words (as seen in the N400), we chose to do it based on quantifying how “semantically dissimilar” each new word was compared to its immediately preceding context. This idea of semantic distance has previously been used in studies of reading-time effects^8^, reading comprehension^19^ and brain imaging of speech processing^19^. Our specific approach was based on the well-known word2vec model^20^, whereby each word in a speech stimulus is converted to a high-dimensional vector (in our case 400 dimensions) that serves as a proxy for that word’s meaning. In particular, words that share common contexts in a very large corpus of text are converted to vectors that are located in close proximity to one another in the high-dimensional space. We then defined the “semantic dissimilarity” of each specific word by comparing (via a Pearson’s correlation) its 400-dimensional vector with the average of the vectors corresponding to all the preceding words in that particular sentence, and then subtracting that correlation from 1. Where a specific word was the first word in a sentence we compared it to the average of all word vectors in the previous sentence, and again subtracted the correlation from 1. This produced a single semantic dissimilarity measure for each word that acts as a representation of the meaning added to a sentence by that word. (Technically this could take any value between 2 and 0, but it tended to be in the range 0.53–1.06). We then created a vector at the same sampling rate as our EEG data (128 Hz) which consisted of time-aligned impulses at the onset of each word that were scaled according to the value of that word’s semantic dissimilarity. Then, by regressing the low-frequency (1–8 Hz) EEG against this vector, we derived a so-called temporal response function (TRF^21^) that describes how these fluctuations in semantic dissimilarity across consecutive words impact upon the neural activity at various time-lags (Fig. 1).

**Figure 1.**
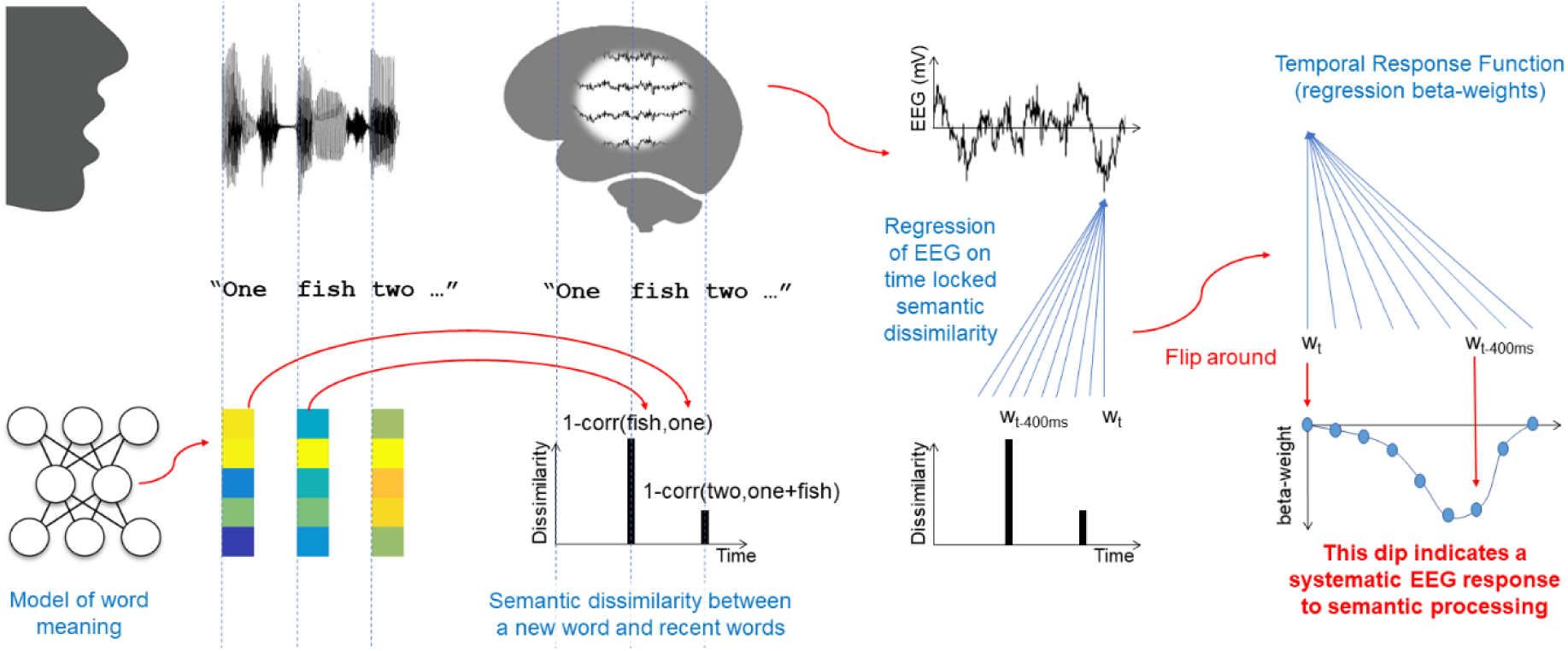
Regression analysis for estimating an electrophysiological correlate of semantic dissimilarity to natural speech. Words from an audiobook are converted to 400-dimensional vectors using the word2vec algorithm^20^ (bottom left). The semantic similarity of each word to its preceding context is then defined by comparing (via a Pearson’s correlation) its 400-dimensional vector with the average of the vectors of all the preceding words in the corresponding sentence. And the “semantic *dis*similarity” of the word is quantified as 1 minus this correlation (bottom middle left). A vector at the same sampling rate as the recorded neural data is then created which consists of time-aligned impulses at the onset of each word that are scaled according to the value of that word’s semantic dissimilarity. The ongoing EEG data is then regressed against this vector to obtain a so-called temporal response function (TRF; right), that describes, via beta-weights, how fluctuations in semantic dissimilarity across words impact upon the EEG at various time-lags ^21^.

### A neural correlate of semantic dissimilarity in natural speech

A TRF averaged over 19 subjects who each listened to ∼60 minutes of an audiobook is shown in Fig. 2A, B. A prominent negativity is apparent over midline parietal scalp at time-lags between 200 and 500 ms (Fig. 2A). Over this time range, this negativity was significantly less than zero across subjects at several parietal scalp electrode sites (Fig. 2B; running one-tailed *t*-test, *P* < 0.05, FDR-corrected). To confirm that this negativity was indeed related to the semantic content of the speech and not just the stimulus acoustics, we repeated the experiment for another 10 subjects who listened to the same audiobook, but in a time-reversed fashion. Conducting the same analysis as before (while taking into account the time-reversed nature of the stimuli) produced TRF responses that showed no evidence of the prominent, late negativity (Fig. 2A, B; running one-tailed *t*-test, *P* > 0.05, FDR-corrected). Indeed, the presence and absence of this negativity for forward and time-reversed speech respectively was evident at the level of several individual subjects who undertook both experiments (Fig. 2C). This pattern of results demonstrates that electrophysiological responses to natural speech, in the form of a late, parietal negativity, reflect the semantic dissimilarity of individual words to their preceding context in a relatively tightly time-locked fashion.

**Figure 2.**
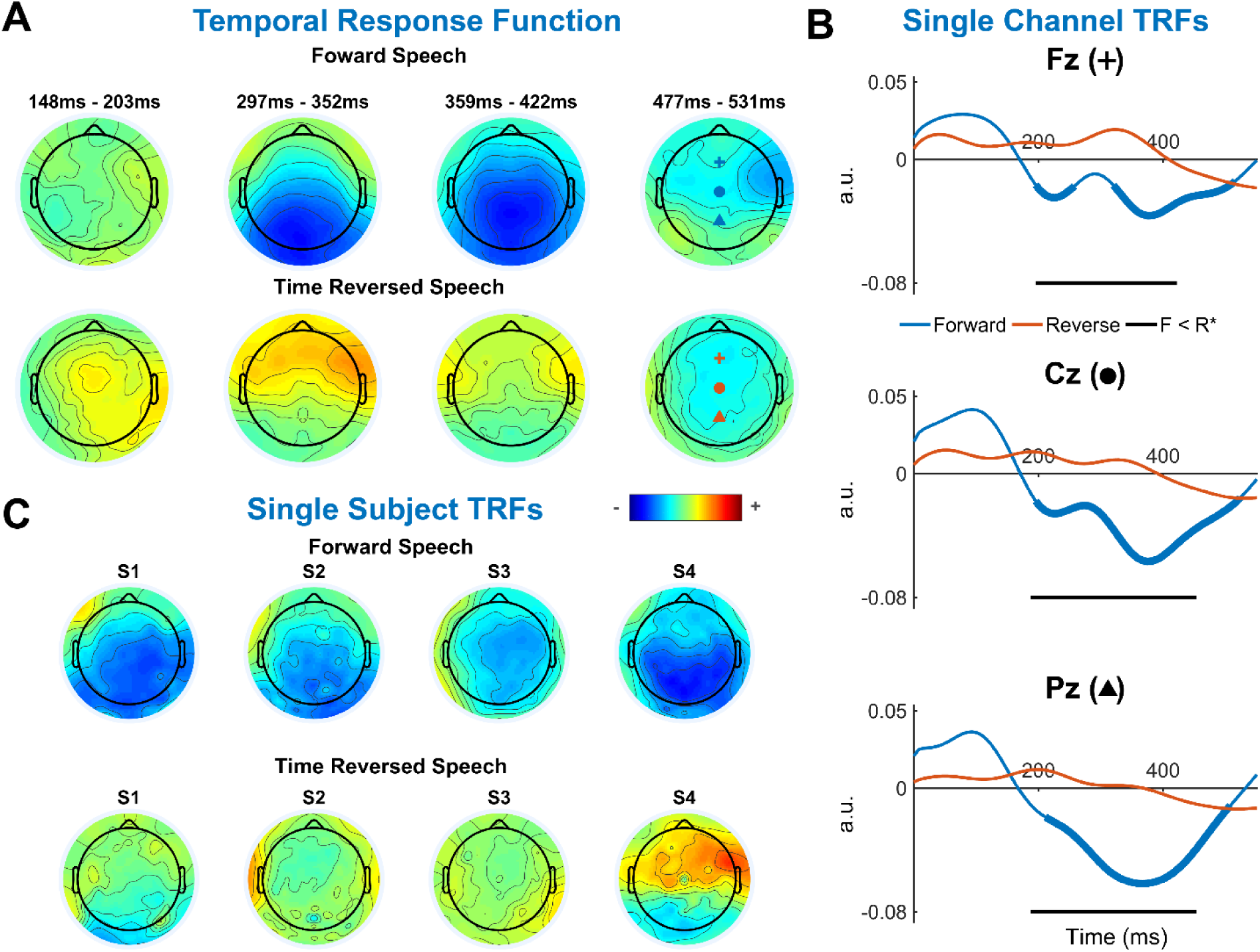
Temporal response functions for natural and time-reversed speech. **A,** Topographic maps of the semantic dissimilarity TRF averaged over all trials and all subjects for natural, forward speech (Top) display a marked centro-parietal negativity between ∼200 and 480 ms. There is no evidence of a similar negativity in the average TRF for time-reversed speech (Bottom). **B,** Grand average TRF waveforms at selected individual channels show the time course of the negativity related to semantic dissimilarity. Thick lines indicate a response that is statistically less than zero across subjects (*P* < 0.05, *t*-test, FDR corrected). And black lines below the waveforms indicate that the TRFs for forward speech are statistically more negative than those for time-reversed speech across subjects (*P* < 0.05, *t*-test, FDR-corrected). **C,** Topographic maps of TRFs averaged over the interval 200–500 ms for selected subjects who took part in both the forward and time-reversed speech experiments. For all four subjects a negativity is apparent for forward speech (albeit with slightly different distributions for each subject) that is absent for the time-reversed speech.

### Neural signatures of semantic dissimilarity depend on intelligibility

The experiments above involved natural speech and its time-reversed counterpart, stimuli that were completely intelligible and completely unintelligible, respectively. To assess how sensitive our semantic dissimilarity TRF might be to gradations of intelligibility we conducted a further experiment involving speech-in-noise. Specifically, we collected EEG data from 21 subjects as they listened to two repetitions of each of fifteen 60-s segments of continuous audio speech which were always mixed with spectrally-matched stationary noise at a signal-to-noise ratio of −9 dB. Based on the well-known fact that visual speech enhances the intelligibility of speech-in-noise^22^, we manipulated intelligibility by allowing subjects to watch a video of the speaker for one of these repetitions. (The presentation order of audio-only, and audio-visual repetitions was randomized across the 15 videos and across subjects). And while the audio-only speech was not completely unintelligible, the presence of the video led to a large, significant improvement in intelligibility as measured by both self-report (*P* = 4 × 10^−5^, Wilcoxon signed rank) and a word detection task (*P* = 7.9 × 10^−5^, Wilcoxon signed rank). This behavioral effect was mirrored by a significant difference in the semantic dissimilarity TRFs between audio-only and audiovisual conditions (Fig. 3A, B). This difference was most pronounced at time-lags between 380 and 600 ms where it showed an effect size of d’ = 0.55. Notably, this was substantially later than the interval for the TRF negativity during clean speech (Fig. 2). Another way to visualize the electrophysiological correlates of improved intelligibility is to assess how well our semantic dissimilarity TRFs can predict unseen EEG responses to natural speech. This kind of forward encoding model-based approach has previously been used for predicting EEG responses to natural speech based on envelope and phonetic representations of speech^18^, as well as fMRI voxel activity based on semantic speech representations^23,24^. Using cross validation to fit a semantic dissimilarity TRF and test it on unseen data produced a significantly better EEG prediction for the audiovisual speech than the audio speech on electrode channels over midline parietal scalp (Fig. 3C; *P* = 0.01, Wilcoxon signed-rank test). And while the EEG predictions based on audio speech were significantly greater than zero – after all, the audio-alone speech was not completely unintelligible – the effect size of adding the visual input on these EEG predictions scores was large (d’ = 0.84 on midline parietal electrode Pz). Overall, this demonstrates that our semantic dissimilarity TRF is sensitive to variations in the intelligibility of acoustically identical speech. Moreover, across subjects in the audiovisual speech condition, there was a significant negative correlation between the self-reported intelligibility ratings (which varied broadly) and the (normalized) amplitude of the TRF negativity averaged over the interval 250 – 500 ms (Fig. 3D; the more intelligible, the larger the negativity; Pearson’s *r* = −0.5, *P* < 0.02). This further highlights the sensitivity of our TRF measure to the intelligibility of natural speech.

### No evidence of contextual semantic processing for unattended speech

Another important test of the behavioral relevance of our semantic dissimilarity TRF would be to determine whether or not it is affected by how much attention a person is paying to natural intelligible speech. Over 60 years ago it was first noted that, when attending to one of two dichotically-presented speech streams, people have a very limited ability to report on the content of the speech in the unattended ear^25^, a phenomenon known as the cocktail party effect. Ever since then, researchers have sought to explain this phenomenon in terms of psychological models^26–28^ and neurophysiological data^15,29–31^. Despite these efforts, the extent to which unattended speech is semantically processed by the brain remains unclear^32,33^. However, given the very marked limitations in the ability of subjects to report on the content of unattended speech, it is reasonable to assert that such speech is not processed to the same depth as attended speech. Thus, we hypothesized that, the negativity in our TRF, as an index of contextual semantic processing, should be markedly reduced in unattended speech.

We recorded EEG from 33 subjects who attended to one of two concurrently and dichotically-presented audiobooks (17 subjects attended to one story and 16 to the other). The experiment was paused after every ∼60s and subjects were tasked with answering multiple-choice questions on both stories. In the same way as the previous experiments, we derived semantic dissimilarity regressors for each of the two stories. And, then we regressed the EEG data against these vectors to produce two TRFs – one for the attended story and one for the unattended story. Consistent with previous studies, the behavioral effect was very strong in this experiment with subjects correctly answering 80% of the questions on the attended story and only 29% of those on the unattended story (chance was 25%). This large behavioral effect was mirrored in differences in the average TRFs across all 33 subjects (Fig. 3E, F). In particular, the TRF corresponding to the attended story showed a clear and prominent negativity over midline parietal scalp, again at a rather long latency in the range 380–600 ms. However, no such negativity was apparent for the unattended speech. While this does not entirely rule out some level of semantic processing in unattended speech – after all, our regressors are based on only one particular computational measure of linguistic processing – it does present strong evidence of a very pronounced reduction in the processing of unattended words relative to their context. The apparently large difference in the magnitude of the negative component between the attended and unattended TRFs over the interval 380–600 ms was supported by statistical testing across subjects (paired *t*-test, *P* = 9.3 × 10^−8^) and showed a very large effect size of d’ = 2.0 on midline parietal electrode Pz. (The effect size over the window 380–550 ms was slightly larger at d’ = 2.25). Again, using cross validation to fit a semantic dissimilarity TRF and then predicting unseen data, produced a significantly better EEG prediction for the attended speech than the unattended speech (Fig. 3G; *P* = 9.36 × 10^−7^, Wilcoxon signed-rank test). And while the EEG predictions based on unattended speech were significantly greater than zero – possibly as a result of weak correlations between the semantic dissimilarity impulses and acoustic energy changes at word onsets – the effect size of attention on these EEG prediction scores was as large as that on the TRFs themselves (d’ = 2.0 on electrode Pz). As for the speech-in-noise experiment above, we wished to investigate the relationship between our TRF measures and behavior more closely. To do this, we checked for across-subject correlations between features of our TRF negativity and subject performance on the attended questions in the cocktail party paradigm. Unlike the audiovisual speech-in-noise where intelligibility varied broadly across subjects, we found no relationship with the amplitude of the TRF and performance on the attended questions. This was not very surprising given that the to-be-attended speech stream was always intelligible. However, we did find that the peak latency of the TRF negativity was significantly negatively correlated with performance on the questions across subjects (*r* = −0.7 *P* =1.952 x 10^−5^). In other words, the earlier a subject’s TRF peak, the better that subject did on the task. We interpret this as evidence that people who can successfully sustain their attention and/or suppress distracting information can more efficiently process the behaviorally relevant speech – or vice versa. This notion of more efficient semantic processing of words in their recent historical context aligns with the well-known link between working memory and cocktail party attention performance^34^.

**Figure 3.**
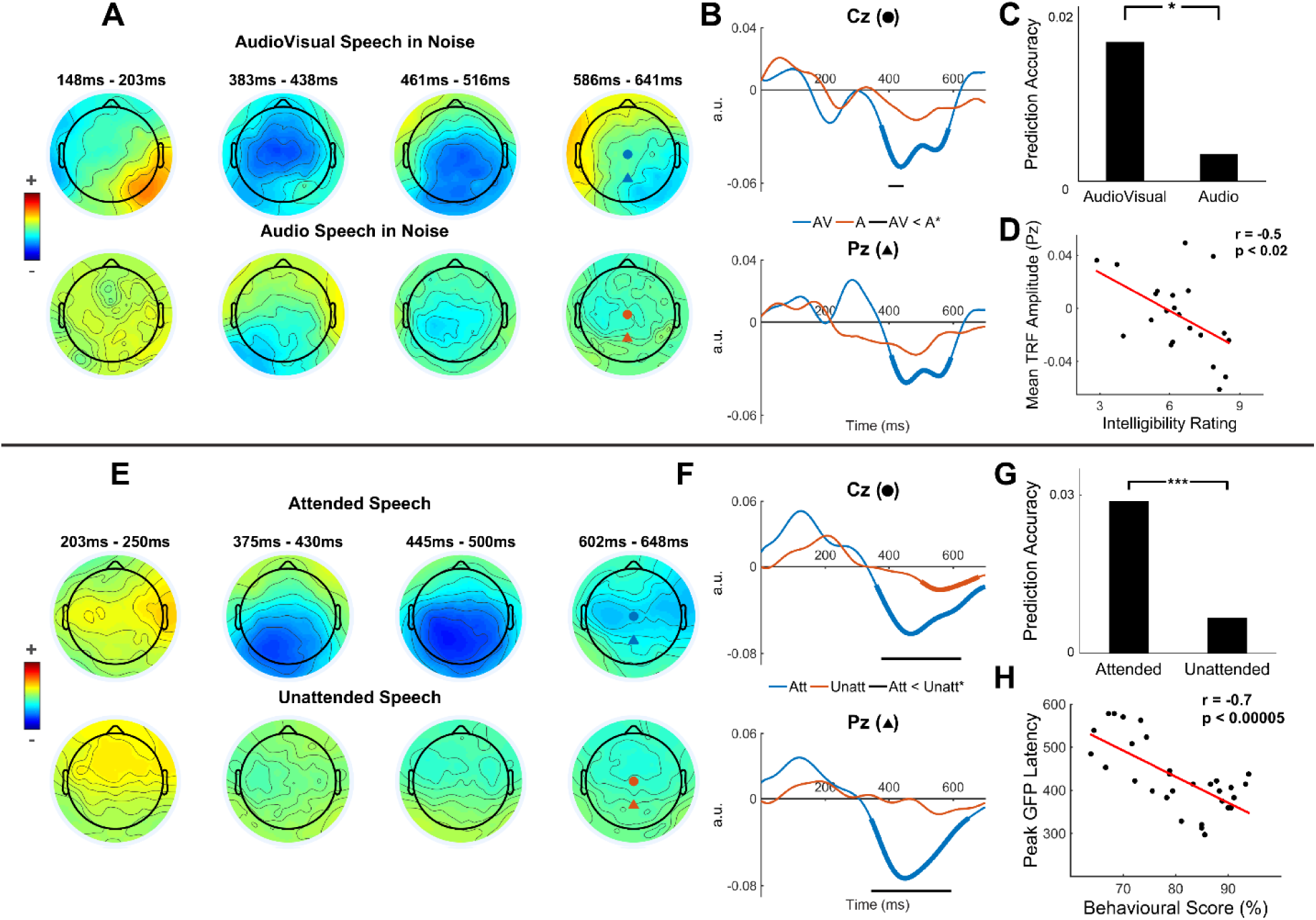
Assessing the effect of comprehension on the electrophysiological index of semantic dissimilarity. **A,** Topographic maps of the semantic dissimilarity TRF averaged over all trials and all subjects for audiovisual speech in −9dB of acoustic background noise display a centro-parietal negativity between ∼400 and 600 ms. This negativity is much reduced in the average TRF for audio-only speech in the same level of background noise, which is much less intelligible. **B,** Grand average TRF waveforms for audiovisual and audio-only speech over two selected midline electrodes. Thick lines indicate a response that is statistically less than zero across subjects (*P* < 0.05, running *t*-test, FDR corrected). And black lines below the waveforms indicate that the TRFs for audiovisual speech are statistically more negative than those for audio-only speech across subjects (*P* < 0.05, running *t*-test, FDR corrected). **C,** A cross-validation procedure was used to predict EEG responses to natural speech using a semantic dissimilarity TRF trained on other data. EEG prediction accuracy for audiovisual speech was significantly greater than that for audio-only speech (*P* < 0.01, *t*-test). **D**, Across subjects, the amplitude of the semantic dissimilarity TRF over midline parietal scalp was significantly correlated with self-reported intelligibility rating of audiovisual speech (*P* < 0.02, Pearson’s correlation). **E,** Topographic maps of the semantic dissimilarity TRF averaged over all trials and all subjects for attended speech in a dichotic cocktail party paradigm display a centro-parietal negativity between ∼300 and 600 ms. This negativity is not apparent in the average TRF for unattended speech. **F,** Grand average TRF waveforms for attended and unattended speech over two selected midline electrodes. Thick lines indicate a response that is statistically less than zero across subjects (*P* < 0.05, running *t*-test, FDR corrected). And black lines below the waveforms indicate that the TRFs for attended speech are statistically more negative than those for unattended speech across subjects (*P* < 0.05, running *t*-test, FDR corrected). **G,** A cross-validation procedure was used to predict EEG responses to natural speech using a semantic dissimilarity TRF trained on other data. EEG prediction accuracy for attended speech was significantly greater than that for unattended speech (*P* < 1 × 10^−6^, *t*-test). **H,** Across subjects, the latency of the peak in the global field power (GFP^35^) of the semantic dissimilarity TRF was significantly negatively correlated with the number of questions answered correctly on the attended speech (*P* < 5 × 10^−5^, Pearson’s correlation).

### Neural correlates of semantic dissimilarity are similar, but not identical to the N400 component

It is noteworthy that the dominant feature of our semantic dissimilarity TRF is a negativity over centro-parietal scalp. This is because the EEG measure that has most strongly been linked with semantic processing – the so-called N400 component – also displays a negative potential over centro-parietal scalp at a latency of around 400 ms. And, while the derivation of the N400 component does not involve the same specific assumptions that underlie our TRF analysis, it is conceivable that differences in semantic dissimilarity accompany the differences in predictability (cloze probability) that drive the development of most N400 stimuli^10^. As such, it is possible that these two measures might reflect, at least partially related processes – or, more specifically, that the N400 contains, as one of several processes, a contribution from the same generators that are driving our TRF. To test this, we recorded EEG data from 9 subjects who undertook a classic N400 experiment and who listened to the audiobook used in our first-mentioned experiment above. For the N400 experiment, subjects read 300 sentences presented word-by-word on a screen, half of which ended with a word that was congruent with the rest of the sentence and half which ended with an incongruent word. N400s were then determined by subtracting the event-related potential to the congruent words from that to the incongruent words. And, using the EEG data recorded during the story, we derived a semantic dissimilarity TRF for each subject as before. Figure 4A, B show that the two responses display somewhat similar timecourses over midline parietal scalp, as well as similar topographical distributions at a latency of 375–425ms. The similarity of the two responses was supported by the fact that the amplitude of the N400 component in the interval 390–450 ms was correlated with the amplitude of the semantic dissimilarity TRF in the interval 330–390 ms across the 9 subjects (Fig. 4C; *r* = 0.751; *P* = 0.0197). The intervals for the two components were chosen based on the distribution of the peak latency for each response type (Fig. 4D). Specifically, these intervals represented the 25^th^ to 75^th^ percentiles of those distributions. Importantly, it should also be noted that these peak latency distributions differed, with the peak latency of the semantic dissimilarity TRF being significantly earlier than that of the N400 (Wilcoxon signed-rank test, *P* = 0.0117; Fig. 4D).

**Figure 4.**
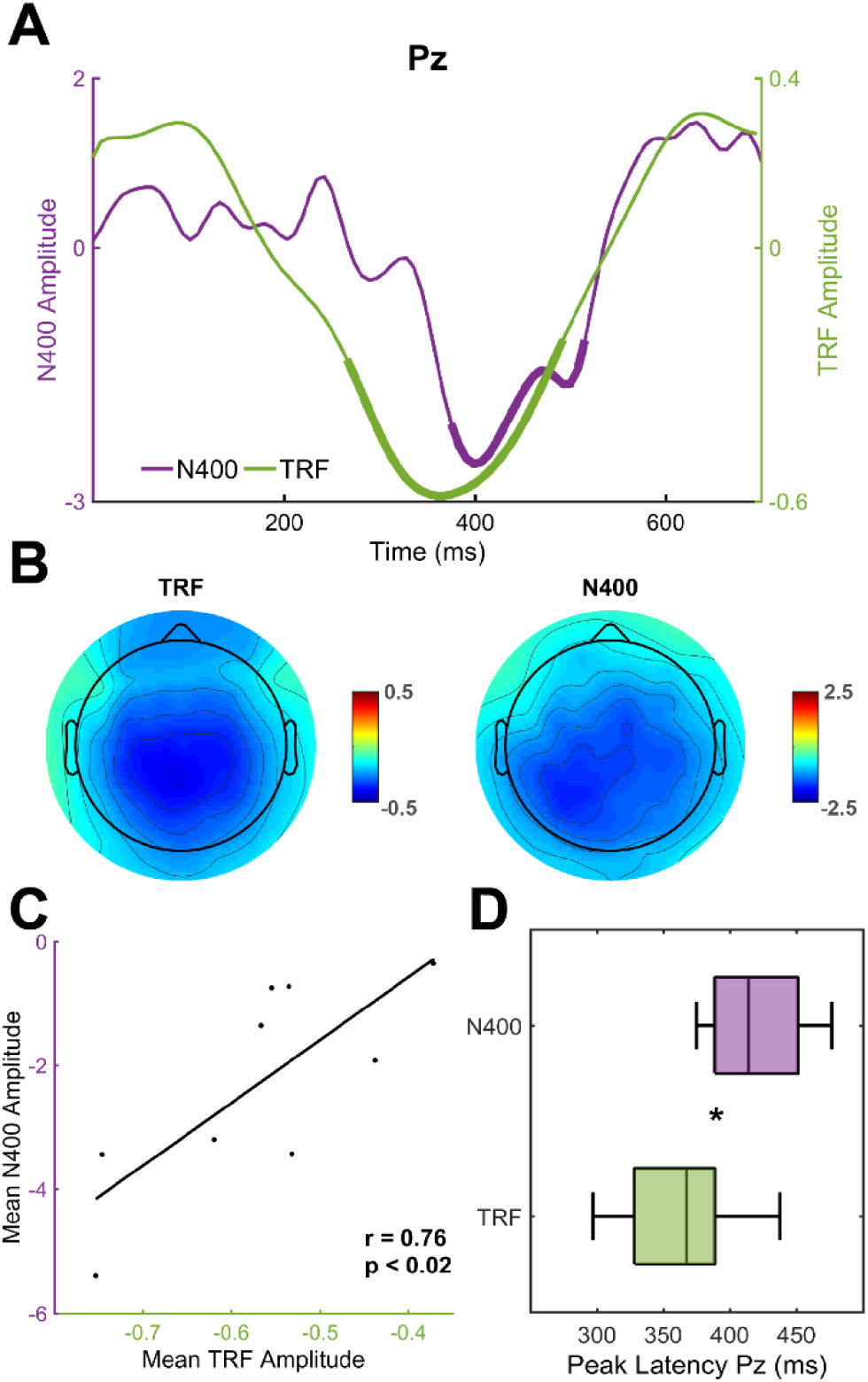
Comparison of semantic dissimilarity TRF and the classic N400 event-related potential component. **A,** Grand average waveforms from a midline-parietal scalp electrode for the classic N400 component (derived by subtracting the average event-related potential to congruent sentence endings from that to incongruent sentence endings) and the semantic dissimilarity TRF. Thick lines indicate a response that is statistically less than zero across subjects (*P* < 0.05, *t*-test). **B,** Topographic maps of both the N400 and the semantic dissimilarity TRF over the interval 375–425 ms. **C,** The amplitudes of the N400 and the semantic dissimilarity TRF are positively correlated across subjects (*P* < 0.02, Pearson’s correlation). **D,** The latency of the peak of the negativity in the semantic dissimilarity TRF is significantly earlier than the peak of the N400 across subjects.

## DISCUSSION

We have shown that, when listening to natural speech, the ongoing dynamics of cortical activity reflect the processing of individual words in the semantic context of the preceding speech in an efficient, time-locked fashion. And we have shown that indices of this processing are robustly affected by whether or not subjects can understand the speech they are hearing and whether or not they are paying attention to that speech. This approach adds an important extra dimension to recent research on the neural tracking of natural speech dynamics by directly linking that tracking to the contextual semantic processing of speech. Further work will be necessary to more fully characterize this online semantic processing. This will include investigating whether or not other types of language knowledge contribute to our measures^19^. It will involve assessing whether unattended speech is processed at a semantic level that depends less upon context than our dissimilarity measure. And it may also include a test of the idea that neural tracking of semantic processing may be even more robust if semantic representation is modeled using a more neurobiologically-motivated approach rather than a model based on word co-occurrence^36^. By incorporating other computational models into the framework we have outlined, we would expect that EEG, ECoG and MEG could be very useful in answering these questions.

It will also be important to more fully examine the similarities and differences between our TRF measure and the N400 component. While the two measures were correlated in terms of amplitude, the TRF negativity peaked significantly earlier than the N400. There are several possible reasons for this latency difference. First, as mentioned above, the assumptions made in deriving the TRF and N400 are not the same; the TRF is based on semantic dissimilarity and the N400 on cloze probability, meaning they may index related, but distinct linguistic processes. Second, it could simply be due to the fact that the TRF is derived using a regression analysis and the N400 using time-locked averaging. We are not convinced of this as an explanation given that our TRF involved representing word meaning as an impulse at word onset, which is a similar assumption to that made when performing averaging of neural data time-locked to word onset. And third, and perhaps most plausibly, the TRF was derived in response to natural, continuous, narrative speech, while the N400 was derived to sentences presented one word at a time. It could be that semantic processing happens more rapidly during natural speech processing, possibly as a result of stronger ongoing predictions in this context^37^, than in the case of individual sentences that may or may not contain incongruent words. The fact that the latency of the TRF negativity was increased for the cocktail party and audiovisual speech-in-noise experiments relative to the first, clean speech experiment, and the fact that working memory has long been linked with speech processing under challenging listening conditions supports this idea of a link between our TRF latency and the efficiency of word processing in the context of preceding words. But, given that the latency of the N400 is typically remarkably constant^10^, and that the amplitude of our TRF negativity was quite strongly correlated with that of the N400, more work will be needed to validate this notion that TRF latency relates to efficient semantic processing. Nonetheless, we suggest here that the interpretation of our TRF can leverage decades of work done on the N400. And that future work using different linguistic models may help to disentangle what might be numerous processes underlying the N400, work that is already ongoing in the domain of reading^19^.

The effects of attention and intelligibility on our electrophysiological measures of semantic similarity closely correlated with the behavioral phenomena in these experiments. And, importantly, these EEG measures were quite robust at the level of individual subjects. This suggests that these indices could be useful in a broad range of basic, applied and clinical research areas. This could include basic research on infant language development, language learning, and the effects of cocktail party attention on speech processing. It could be helpful in arbitrating between computational models of natural speech processing. It could be of benefit to researchers studying language impairments in different clinical cohorts, in clinical testing of disorders of consciousness, and, given that it relies on relating the processing of words to their recent context, as an assay in people at risk for cognitive and memory decline.

## METHODS

All procedures were undertaken in accordance with the Declaration of Helsinki and were approved by the Ethics Committees of the School of Psychology at Trinity College Dublin, and the Health Sciences Faculty at Trinity College Dublin.

### Subjects

All subjects were native English speakers, and reported normal hearing, normal or corrected-to-normal vision, and no history neurological disease. 19 subjects (13 Male) aged between 19 and 38 years participated in the first experiment involving listening to a single audiobook. Of these 19 subjects, 9 also participated in the N400 experiment. And ten subjects (7 male) aged between 21 and 32 years participated in the experiment involving the time-reversed audiobook (5 of these subjects participated in the first experiment above). 34 subjects (28 male) with a mean ± SD age of 27.3 ± 3.2 years participated in the cocktail party attention experiment, but data from one subject was not included in the analysis as the recordings from their mastoid electrodes were of poor quality. And 21 subjects (6 female) aged between 21 and 35 years participated in the multisensory speech experiment. Much of the data from these experiments has previously been published in studies examining how EEG tracks the envelope and phonetic content of speech^18,38,39^.

### Data acquisition and pre-processing

For all experiments, 128-channel EEG data (plus two mastoid channels) were acquired at a rate of 512 Hz using an ActiveTwo system (BioSemi). Triggers indicating the start of each trial were sent by the stimulus presentation computer and included in the EEG recordings to ensure synchronization. Offline, the data were band-pass filtered between 1 and 8 Hz, downsampled to 128 Hz, and re-referenced to the average of the mastoid channels in MATLAB. To identify channels with excessive noise, the time series were visually inspected and the SD of each channel was compared with that of the surrounding channels. Channels contaminated by noise were recalculated by spline interpolating the surrounding clean channels in EEGLAB ^40^.

### Stimuli and Procedures

In the first experiment, subjects undertook 20 trials, each of the same length (just under 180 seconds), where they were presented with a professional audio-book version of a popular mid-20th century American work of fiction written in an economical and understated style and read by a single male American speaker. The trials preserved the storyline, with neither repetitions nor discontinuities. The average speech rate was ∼210 words/min. Similarly, the second experiment involved the presentation of the same trials in the same order, but with each of the 28 speech segments played in reverse. All stimuli were presented monophonically at a sampling rate of 44,100 Hz using Sennheiser HD650 headphones and Presentation software from Neurobehavioral Systems (http://www.neurobs.com). Testing was carried out in a dark room and subjects were instructed to maintain visual fixation for the duration of each trial on a crosshair centered on the screen, and to minimize eye blinking and all other motor activities.

In the cocktail party experiment, subjects undertook 30 trials, each of ∼1 min in length, where they were presented with 2 classic works of fiction: one to the left ear, and the other to the right ear. Each story was read by a different male speaker. Subjects were divided into 2 groups of 20 with each group instructed to attend to the story in either the left or right ear throughout all 30 trials. After each trial, subjects were required to answer between 4 and 6 multiple-choice questions on both stories. Each question had 4 possible answers. We used a between-subjects design as we wanted each subject to follow just one story to make the experiment as natural as possible and because we wished to avoid any repeated presentation of stimuli. For both stories, each trial began where the story ended on the previous trial. Stimulus amplitudes in each audio stream within each trial were normalized to have the same root mean squared (RMS) intensity. In order to minimize the possibility of the unattended stream capturing the subjects’ attention during silent periods in the attended stream, silent gaps exceeding 0.5 s were truncated to 0.5 s in duration. Stimuli were presented using Sennheiser HD650 headphones and Presentation software from Neurobehavioral Systems (http://www.neurobs.com). Subjects were instructed to maintain visual fixation for the duration of each trial on a crosshair centered on the screen, and to minimize eye blinking and all other motor activities.

For the multisensory experiment, the stimuli were drawn from a set of videos that consisted of a male speaking American English in a conversational-like manner. Fifteen 60-s videos were rendered into 1280 x 720-pixel movies at 30 frames/s and exported in audio-only (A), visual-only (V), and AV format in VideoPad Video Editor (NCH Software). The soundtracks were sampled at 48 kHz, underwent dynamic range compression, and were matched in intensity (as measured by root mean square; see ^38^), and were mixed with spectrally matched stationary noise to ensure consistent masking across stimuli ^41,42^. The noise stimuli were generated in MATLAB (The MathWorks) using a 50th-order forward linear predictive model estimated from the original speech recording. Prediction order was calculated based on the sampling rate of the soundtracks ^43^.

### Computational model and regression

Semantic vectors for words were derived using the state-of-the-art word2vec algorithm ^20^. The “continuous bag of words” implementation ^44^ was selected, because this was built on British English corpora (ukWaC, the English Wikipedia and the British National Corpus combined) which is both large and probably more reflective of the language exposure of the participants (in Dublin) than US corpora. In addition, word vectors are freely downloadable (see ^44^). Word2vec embodies the “distributional hypothesis” that words with similar meaning occur in similar contexts in an artificial neural network approach. Practically, the approach involves sliding a fixed window of words (11 in this case, however this is a parameter set by the experimenter) over a text corpus and training a neural network to predict the word in the center of that window. Word identity (as opposed to semantics) is uniquely encoded as a single bit set to one in a long vector of zeros (vector length is the number of words in the vocabulary). These long vectors form the basis of the input and output to the neural network. The input corresponds to the sum of the 10 word vectors in the window, the output is the central word. Because word order is lost in this summation, the input is analogous to an unordered bag of words. The network contains an internal hidden layer of 400 dimensions (400 is also a parameter set by the experimenter). The hidden layer is fully connected to the input and output. It is in fact the weights on the connections between the input and hidden layer that are ultimately harvested to form the semantic model (the weights are a number-of-words in the vocabulary by 400 floating point matrix) and the remainder of the network is discarded. Weights are initially set as random, but are subsequently optimized so as to reduce error between predicted and target output. Intuitively, because words that frequently appear together in the same context window also predict similar central words, weights on these words are tuned to similar internal representations reflecting common contexts. For more details on the training procedure see ^44^ and ^20^.

Having obtained a vector for each word, we then quantified how semantically dissimilar each particular word was to the preceding words in the corresponding sentence. We did this by calculating a Pearson’s correlation between the word’s 400-dimensional vector and the average of the vectors corresponding to all the preceding words in that particular sentence, and subtracting this correlation from 1. (Where a specific word was the first word in a sentence, we calculated the correlation between the word and the average of all word vectors in the previous sentence, before, again, subtracting that correlation from 1). This produced a single semantic dissimilarity measure for each word with a value between 2 and 0. We then created a “semantic dissimilarity vector” at the same sampling rate as our EEG data (128 Hz) which consisted of time-aligned impulses at the onset of each word that were scaled according to the value of that word’s semantic dissimilarity. The word onset times were determined by performing forced alignment of the speech files and the corresponding textual orthographical transcription using the Prosodylab-Aligner ^45^.

The method used here to analyze the mapping between the semantic dissimilarity vector and the recorded EEG data is commonly known as a temporal response function (TRF). A TRF can be interpreted as a filter that describes the brain’s linear transformation of a stimulus feature, S(*t*), to the continuous neural response R(t), i.e., R(*t*) = TRF*S(*t*) where ‘*’ represents the convolution operator. The TRFs were calculated by performing regularized linear regression between our stimulus variables and our EEG. Specifically, we performed ridge regression wherein a parameter (lambda) is set to control overfitting (see 21 for a detailed description of this step).

In previous work, we have attempt to cast our TRF functions with μV as their unit of measure. However, this relies on a decision to normalize the input stimulus values between some limits and, as such, has been somewhat arbitrary. In the present work, and in line with previous work from other groups, the EEG data on each channel was z-scored prior to estimating the TRF, meaning that the TRFs are ultimately presented in arbitrary units. The colors in the TRF topographic plots can be interpreted as follows: red at a particular latency indicates that, at that poststimulus lag, the EEG voltage is driven in a positive direction by a unit change in semantic dissimilarity. And blue means the EEG voltage at that poststimulus lag is driven negative by a similar change. Thus, given the same normalization strategy for the various speech stimuli used in this study, the TRF responses can be compared in terms of their amplitudes, despite their description in terms of arbitrary units.

## Acknowledgements

This study was supported by the Irish Research Council through the Government of Ireland Postgraduate Scholarship scheme (M.P.B. & G.D.L.) and a Career Development Award from Science Foundation Ireland (E.C.L.).

## Author Contributions

E.C.L., G.D.L., and A.J.A. conceived of the experiment. M.B, G.D.L., and M.J.C. collected data. M.B., G.D.L., and A.J.A. analyzed the data. E.C.L. wrote the first draft of the manuscript. M.B., G.D.L. and A.J.A edited the manuscript.

## References

1 Crystal, T. H. & House, A. S. Articulation rate and the duration of syllables and stress groups in connected speech. The Journal of the Acoustical Society of America 88, 101–112 (1990).

2 Liberman, A. M., Cooper, F. S., Shankweiler, D. P. & Studdert-Kennedy, M. Perception of the speech code. Psychol. Rev. 74, 431 (1967).

3 Simpson, G. B. Understanding word and sentence. Vol. 77 (Elsevier, 1991).

4 Marslen-Wilson, W. Linguistic structure and speech shadowing at very short latencies. Nature (1973).

5 Tanenhaus, M. K., Spivey-Knowlton, M. J., Eberhard, K. M. & Sedivy, J. C. Integration of visual and linguistic information in spoken language comprehension. Science, 1632–1634 (1995).

6 Levy, R. Expectation-based syntactic comprehension. Cognition 106, 1126–1177 (2008).

7 Landauer, T. K. & Dumais, S. T. A solution to Plato’s problem: The latent semantic analysis theory of acquisition, induction, and representation of knowledge. Psychol. Rev. 104, 211 (1997).

8 Pynte, J., New, B. & Kennedy, A. On-line contextual influences during reading normal text: A multiple-regression analysis. Vision Res. 48, 2172–2183 (2008).

9 Mitchell, J., Lapata, M., Demberg, V. & Keller, F. in Proceedings of the 48th Annual Meeting of the Association for Computational Linguistics. 196–206 (Association for Computational Linguistics).

10 Kutas, M. & Federmeier, K. D. Thirty years and counting: Finding meaning in the N400 component of the event related brain potential (ERP). Annu. Rev. Psychol. 62, 621 (2011).

11 Ahissar, E. et al. Speech comprehension is correlated with temporal response patterns recorded from auditory cortex. Proceedings of the National Academy of Sciences 98, 13367–13372 (2001).

12 Lalor, E. C. & Foxe, J. J. Neural responses to uninterrupted natural speech can be extracted with precise temporal resolution. Eur. J. Neurosci. 31, 189–193, doi:10.1111/j.1460-9568.2009.07055.x (2010).

13 Luo, H. & Poeppel, D. Phase patterns of neuronal responses reliably discriminate speech in human auditory cortex. Neuron 54, 1001–1010 (2007).

14 Ding, N. & Simon, J. Z. Emergence of neural encoding of auditory objects while listening to competing speakers. Proceedings of the National Academy of Sciences 109, 11854–11859 (2012).

15 Power, A. J., Foxe, J. J., Forde, E. J., Reilly, R. B. & Lalor, E. C. At what time is the cocktail party? A late locus of selective attention to natural speech. Eur. J. Neurosci. 35, 1497–1503, doi:10.1111/j.1460-9568.2012.08060.x (2012).

16 Peelle, J. E., Gross, J. & Davis, M. H. Phase-locked responses to speech in human auditory cortex are enhanced during comprehension. Cerebral cortex 23, 1378–1387 (2013).

17 Ding, N., Melloni, L., Zhang, H., Tian, X. & Poeppel, D. Cortical tracking of hierarchical linguistic structures in connected speech. Nat. Neurosci. 19, 158–164, doi:10.1038/nn.4186 (2016).

18 Di Liberto, G. M., O’Sullivan, J. A. & Lalor, E. C. Low-Frequency Cortical Entrainment to Speech Reflects Phoneme-Level Processing. Curr. Biol. 25, 2457–2465 (2015).

19 Frank, S. L. & Willems, R. M. Word predictability and semantic similarity show distinct patterns of brain activity during language comprehension. Language, Cognition and Neuroscience, 1–12 (2017).

20 Mikolov, T., Chen, K., Corrado, G. & Dean, J. Efficient estimation of word representations in vector space. arXiv preprint arXiv:1301.3781 (2013).

21 Crosse, M. J., Di Liberto, G. M., Bednar, A. & Lalor, E. C. The multivariate temporal response function (mTRF) toolbox: a MATLAB toolbox for relating neural signals to continuous stimuli. Front. Hum. Neurosci. 10 (2016).

22 Sumby, W. H. & Pollack, I. Visual contribution to speech intelligibility in noise. J. Acoust. Soc. Am. 26, 212–215, doi:10.1121/1.1907309 (1954).

23 de Heer, W. A., Huth, A. G., Griffiths, T. L., Gallant, J. L. & Theunissen, F. E. The hierarchical cortical organization of human speech processing. J. Neurosci., 3267–3216 (2017).

24 Huth, A. G., de Heer, W. A., Griffiths, T. L., Theunissen, F. E. & Gallant, J. L. Natural speech reveals the semantic maps that tile human cerebral cortex. Nature 532, 453–458 (2016).

25 Cherry, E. C. Some experiments on the recognition of speech, with one and with two ears. The Journal of the acoustical society of America 25, 975–979 (1953).

26 Broadbent, D. E. Perception and communication. (Pergamon Press, 1958).

27 Deutsch, J. A. & Deutsch, D. Attention: some theoretical considerations. Psychol. Rev. 70, 80 (1963).

28 Treisman, A. M. Verbal cues, language, and meaning in selective attention. The American journal of psychology, 206–219 (1964).

29 Mesgarani, N. & Chang, E. F. Selective cortical representation of attended speaker in multi-talker speech perception. Nature 485, 233–236 (2012).

30 Teder, W., Kujala, T. & Näätänen, R. Selection of speech messages in free-field listening. Neuroreport: An International Journal for the Rapid Communication of Research in Neuroscience (1993).

31 Zion-Golumbic, E. M. et al. Mechanisms Underlying Selective Neuronal Tracking of Attended Speech at a “Cocktail Party” Neuron 77, 980–991, doi:10.1016/j.neuron.2012.12.037 (2013).

32 Aydelott, J., Jamaluddin, Z. & Nixon Pearce, S. Semantic processing of unattended speech in dichotic listening. The Journal of the Acoustical Society of America 138, 964–975 (2015).

33 Lachter, J., Forster, K. I. & Ruthruff, E. Forty-five years after Broadbent (1958): still no identification without attention. Psychol. Rev. 111, 880 (2004).

34 Conway, A. R., Cowan, N. & Bunting, M. F. The cocktail party phenomenon revisited: The importance of working memory capacity. Psychonomic bulletin & review 8, 331–335 (2001).

35 Lehmann, D. & Skrandies, W. Reference-free identification of components of checkerboard-evoked multichannel potential fields. Electroencephalogr. Clin. Neurophysiol. 48, 609–621, doi:10.1016/0013-4694(80)90419-8 (1980).

36 Anderson, A. J. et al. Predicting neural activity patterns associated with sentences using a neurobiologically motivated model of semantic representation. Cerebral Cortex (2016).

37 Kuperberg, G. R. & Jaeger, T. F. What do we mean by prediction in language comprehension? Language, cognition and neuroscience 31, 32–59 (2016).

38 Crosse, M. J., Di Liberto, G. M. & Lalor, E. C. Eye Can Hear Clearly Now: Inverse Effectiveness in Natural Audiovisual Speech Processing Relies on Long-Term Crossmodal Temporal Integration. J. Neurosci. 36, 9888–9895, doi:10.1523/jneurosci.1396-16.2016 (2016).

39 O’Sullivan, J. A. et al. Attentional selection in a cocktail party environment can be decoded from single-trial EEG. Cerebral Cortex 25, 1697–1706 (2015).

40 Delorme, A. & Makeig, S. EEGLAB: an open source toolbox for analysis of single-trial EEG dynamics including independent component analysis. J. Neurosci. Methods 134, 9–21, doi:10.1016/j.jneumeth.2003.10.009 (2004).

41 Ding, N., Chatterjee, M. & Simon, J. Z. Robust cortical entrainment to the speech envelope relies on the spectro-temporal fine structure. Neuroimage 88, 41–46 (2014).

42 Ding, N. & Simon, J. Z. Adaptive Temporal Encoding Leads to a Background-Insensitive Cortical Representation of Speech. J. Neurosci. 33, 5728–5735, doi:10.1523/jneurosci.5297-12.2013 (2013).

43 Parsons, T. W. Voice and speech processing. (McGraw-Hill College, 1987).

44 Baroni, M., Dinu, G. & Kruszewski, G. in *ACL (1).* 238–247.

45 Gorman, K., Howell, J. & Wagner, M. Prosodylab-aligner: A tool for forced alignment of laboratory speech. Can. Acoustics 39, 192–193 (2011).

